# Comparative analysis of maternal gene expression patterns unravels evolutionary signatures across reproductive modes

**DOI:** 10.1101/2023.11.01.565082

**Authors:** Ferenc Kagan, Andreas Hejnol

## Abstract

**Background:** Maternal genes have a pivotal role in regulating metazoan early development. As such their functions have been extensively studied since the dawn of developmental biology. The temporal and spatial dynamics of their transcripts have been thoroughly described in model organisms and their functions have been undergoing heavy investigations. Yet, less is known about the evolutionary changes shaping their presence within diverse oocytes. Due to their unique maternal inheritance pattern, a high degree is predicted to be present when it comes to their expression. Insofar only limited and conflicting results have emerged around it.

**Results:** Here we set out to elucidate which evolutionary changes could be detected in the maternal gene expression patterns using phylogenetic comparative methods on RNAseq data from 43 species. Using normalized gene expression values and fold change information throughout early development we set out to find the best fitting evolutionary model. Through modeling we find evidence supporting both the high degree of divergence and constraint on gene expression values, together with their temporal dynamics. Furthermore, we find that maternal gene expression alone can be used to explain the reproductive modes of different species.

**Conclusions:** Together, these results suggest a highly dynamic evolutionary landscape of maternal gene expression. We also propose a possible functional dichotomy of maternal genes which is influenced by the reproductive strategy undertaken by examined species.

## 1. Introduction

Early metazoan development is characterized by fast cell divisions, which do not allow *de novo* transcription. Yet, the development still commences due to a set of predetermined factors present already in the oocytes. These factors are the RNA and protein products of maternal genes (Harvey, 1936; Stroband et al., 1992).

Among various functions, some better-known gene products of maternal genes are responsible for the maintenance of the chromatin state (Golding et al., 2011; Hirasawa et al., 2008) and in parallel the activation of the silenced zygotic genome (Bultman et al., 2006; Gu et al., 2011; Pan & Schultz, 2011). Evidence also suggests roles in cell adhesion (de Vries et al., 2004; Larue et al., 1994), inhibition of polyspermy (Burkart et al., 2012), and patterning of the early embryo (Lehmann & Nusslein-Volhard, 1991).Maternal genes exhibit autoregulation through molecular complexes responsible for breaking down maternal transcripts (Lykke-Andersen et al., 2008; Tsukamoto et al., 2008). Together with activation of *de novo* transcribed factors, the maternal genes become degraded gradually from the early embryo through a period that was defined as the maternal-to-zygotic transition (MZT) (Vastenhouw et al., 2019). Generally, the MZT can be separated into two major regulatory phases. The first phase can be characterized by post-transcriptional (Stoeckius et al., 2014; Thomsen et al., 2010) and post-translational (Krauchunas et al., 2012) regulatory events. I.e. it has been shown previously that maternal transcripts possess longer 3’ untranslated regions (3’-UTRs), an observation attributed to post-transcriptional regulation converging on *cis* motifs found in these regions (Shen-Orr et al., 2010). During the second regulatory phase, the zygotic genome is activated, therefore the major regulatory events will shift towards a transcriptional one (De Iaco et al., 2017; Lee et al., 2014). The transition from a maternal control to a zygotic control happens on a large scale, as maternally provided transcripts can take up to three-quarters of the zygotic transcriptome (Thomsen et al., 2010). Despite its complexity and magnitude, the MZT is a highly conserved transition present in all metazoan species (Vastenhouw et al., 2019) and even some plant species (Zhao et al., 2022). Several functions have been proposed for the MZT, such as acting as an internal clock (Tadros & Lipshitz, 2009), a major reprogramming event (Lee et al., 2014), or the degradation of oogenesis-specific transcripts (Pan et al., 2005).

Maternal genes fall under the umbrella term of maternal effects, a term originally coined by Timothy A. Mousseau and Charles W. Fox (Mousseau & Fox, 1998, page 5). According to their model, the genes falling under the maternal effect will show higher evolutionary divergences compared to the genes which have no such effects present. Furthermore, this weakened selection is further enhanced by a generational uncoupling, meaning that the population under selection will not respond to selection in the same generation, instead, there will be a lag of one generation (Kirkpatrick & Lande, 1989). Empirical evidence has emerged throughout the years which suggests that the theoretical predictions might hold true (Demuth & Wade, 2007; Cruickshank & Wade, 2008).

Comparative studies have had a strong resurgence in the past few decades, due to the constant development of robust statistical analyses and the implementation of these into user-friendly libraries (see f.ex. Pennell et al., 2014; Revell, 2012). In parallel, the development of high-throughput data generation has seen an unprecedented expansion. Together, the two opened the avenue for research on the evolutionary forces shaping molecular phenomena through comparative analyses. Indeed, recent studies focused on gene expression datasets in a comparative framework to decipher evolutionary forces shaping the transcriptomes. Focus has been given to mammalian organ gene expression evolution (Brawand et al., 2011; Cardoso-Moreira et al., 2019; Fukushima & Pollock, 2020; Guschanski et al., 2017). Some studies ventured into elucidating narrower evolutionary questions, such as siphonophore transcriptome evolution (Munro et al., 2022) or the ovary specific gene expression in Drosophiliids (Church et al., 2023). All these studies share in common the use of phylogenetic comparative methods, an approach proposed by Felsenstein in his seminal article (Felsenstein, 1985). There, Felsentsein argued against the use of standard statistical tools within comparative biology as species share a common ancestry. Standard statistical tools are ill-equipped to deal with such covariation, therefore he proposed an alternative approach in the form of independent contrasts (Felsenstein, 1985). Since then various other approaches have been developed to deal with biological data in a comparative framework (Freckleton et al., 2002; Pagel, 1997; Pagel, 1999; Rohlfs & Nielsen, 2015).

Here we set out to elucidate the evolutionary patterns shaping maternal gene expression evolution using phylogenetic comparative methods. Results from theoretical approaches (Mousseau & Fox, 1998) and sequence based methods (Cruickshank & Wade, 2008; Demuth & Wade, 2007) suggest a highly dynamic evolutionary landscape of maternal genes. On contrary, more recent studies on this topic (Atallah & Lott, 2018a, 2018b) point towards the maternal transcriptome being conserved. However, these studies are limited in species included and omitted the use of phylogenetic comparative methods. To determine which scenario is more plausible we set out to study the maternal transcriptome evolution in a phylogenetic comparative framework with an expanded set of species included in the analysis. Our results suggest the presence of signals for divergence and constraint being simultaneously present in the maternal transcriptome.

## 2. Material and methods

### 2.1 Quantification and differential gene expression analysis

Following data retrieval, preprocessing, and assemblies (see Suppl. Methods 1.4), the quantification step was performed with the salmon’s pseudoaligner algorithm (Patro et al., 2017). To ensure the highest quality of the alignment step we followed the suggestions of the developers (Suppl. methods). The alignments rates were inspected before downstream use and low-quality samples were discarded.

For differential gene expression analysis custom R scripts (R Core Team, 2022) were written which utilized widely used libraries for downstream analyses (Suppl. Methods and GitHub repository). Here we define a maternal transcript as any transcript with a transcript per million (TPM) value above 2 within the oocyte stages of the sampled species. This cutoff was chosen following previous results (Wagner et al., 2013). For DGE a standard pipeline of tximport (Soneson et al., 2015) and DESeq2 (Love et al., 2014) pipeline was used. Not all species have a reported timeframe for MZT, therefore contrasting points had to be determined. Euclidean distances were quantified across the variance-stabilized samples. Compared to the oocyte stages the first developmental stage with the highest distances was searched by pairwise comparisons of the samples. This was confirmed by clustering the samples based on Euclidean distances. The heatmap provided visual information for shifts in the transcriptome and in conjunction with the known developmental stages the earliest major transcriptional shift was set as an anchoring point for the MZT. To account for unknown variables during data collection a surrogate variable analysis was performed using the sva (Leek et al., 2012), and these variables were incorporated in the design formula during differential gene expression analysis. Where variables apart from the developmental stages were known the design formulas were set up with these accounted for.

Following the above preparations, the dynamics of maternal genes were determined with cutoff values for adjusted p-values of 0.05 and log2 fold change ± 2. To validate DGE results *in-situ* hybridization-based categorizations from the Fly-FISH database (Lécuyer et al., 2007, Wilk et al., 2016) were retrieved and compared to the *Drosophila melanogaster* list of degraded genes.

Information on the gene architectural features of maternal genes is scarce to our knowledge (Heyn et al., 2014), therefore having such a dataset available would be valuable. We set out to inspect the general architectural characteristics. Furthermore, their stability is highly regulated through the binding of regulator proteins to the 3’ untranslated region (3’-UTR), therefore having an overview of such features could provide valuable information (Mishima & Tomari, 2016). If there were gene models with the right information (i.e. annotated genome), we took the lengths of different features for maternal genes that weren’t degraded. We also did this for maternal genes that were degraded and for genes that weren’t expressed maternally. These lengths were then directly compared between downregulated and non-downregulated maternal genes by amalgamating the information from each species.

### 2.2 Functional enrichment

Functional enrichment analysis was done in R programming language environment (R Core Team, 2022). Functional annotations were either imported from available genomic resources or assigned *de novo*. For the latter two strategies were used: the transcriptome assembly provided homology information was used for inferring gene ontology annotations or a web-service tool Pannzer2 assigned high probability annotations. For the enrichment itself the enricher and enrichGO functions were utilized (Yu et al., 2012). The former was used in cases with custom gene ontological annotation databases built *de novo*, the latter for available annotations. If custom annotations were provided to enricher() as a background set all GO annotations retrieved for all genes per each species were used. All ontological categories were tested and considered enriched with a cut-off value of < 0.05 for the adjusted p-values. Both categories of maternal genes were tested this way separately, ordering of the genes was done by the TPM values for the maternal genes.

Functional enrichments were performed on orthogroups also. In these cases, orthogroups were first annotated using the UniProt database (Bateman et al., 2021). Following the annotation, all GO terms attached to the most probable annotation for each orthogroup were retrieved using UniProtR (Soudy et al., 2020). As background set, all GO terms retrieved for all orthogroups were selected. Enrichment was performed using the hypergeometric test mentioned above. The terms below adjusted p-values of 0.05 were selected.

### 2.3 Orthology mapping

To determine orthology relationships we kept only the longest versions of genes from the translated CDSs and used OrthoFinder2 (Emms & Kelly, 2015, 2019) to map them out. We performed the sequence alignments using the diamond’s ultra-sensitive mode (Buchfink et al., 2014), followed by clustering with the default inflation parameter, and gene trees were estimated in a multiple sequence alignment mode using MAFFT (Katoh & Standley, 2013). We used a species tree retrieved from the Open Tree of Life database with the rotl (Michonneau et al., 2016; Open Tree Of Life et al., 2019) during orthology assignments as a starting tree.

In order to use phylogenetic comparative methods, a dated species tree was required. We calibrated the species tree generated by OrthoFinder using geiger’s congruify approach (Eastman et al., 2013). We accessed internal branching event timings from the TimeTree database (Kumar et al., 2017). For scaling the branch lengths, we used TreePL (Smith & O’Meara, 2012). We ran the scaling process in three stages: first, an initial optimization run; second, a run testing different smoothing parameter values; and finally, a run with all parameters set to optimal values to perform the dating.

### 2.4 Phylogenetic dataset assimilation

To compare gene expressions across different species, a data matrix was created by combining orthology relationships with TPM values. The raw TPM values were first normalized within each species using edgeR’s TMM method (Robinson et al., 2009), then across species using a recently proposed method (Munro et al., 2022). Following this, replicates were collapsed into a single value by calculating their means and this single value was assigned to each orthogroup for given species. If multiple genes were assigned to one orthogroup, then these paralogs were summarized into a single value represented by their mean. The values were log-transformed with a pseudocount of 0.01 To account for batch effects, available metadata was fed into limma’s removeBatchEffect function (Ritchie et al., 2015).

### 2.5 Evolutionary modeling

Before applying phylogenetic aware models to the maternal gene expression dataset we had to ensure that the use of phylogenetic models is indeed justified. For this, we measured the phylogenetic signal present in each orthogroup utilizing Blomberg’s approach (Blomberg et al., 2003). Significant values from the randomization test indicate that a phylogenetic signal is present, whilst the K metric indicates the departure from a Brownian motion assumption. A K value of 1 specifies the expected variation of the trait under a Brownian motion model. K values higher than 1 suggest that relatives resemble each other more than expected, whilst K values lower than 1 suggest more dissimilarity among relatives than expected. A driving force for the former could be selection, whilst for the latter homoplasy.

A step for classifying reproductive modes was inserted as a first step in order to use multi-regime evolutionary models. The reproductive mode for each species was determined using the classification of Thierry Lodé (Lodé, 2012). Species with placenta and giving live birth according to this classification follow the hemotrophic viviparity mode of reproduction (such as for example Homo sapiens, and Mus musculus). Oviparitic species can be characterized by internal fertilization and the embryos are supplied with high quantities of yolk (such as for example *Caenorhabditis elegan*s, and *Drosophila melanogaster*). Finally, ovuliparitic species utilize external fertilization with a moderate amount of yolk supplied with the oocytes. The fourth category, the histotrophic viviparity was not included in the analyzed dataset as to our knowledge there are no available datasets sufficing our criteria of inclusion. Included modes of reproduction have been mapped on the species tree using phytool’s make.simmap (Revell, 2012) function. This was necessary for fitting evolutionary models that enable different rates across the mapped character states (reproductive modes in this case).

Before the model fitting step in each orthogroup the species tree was pruned for the tips which have maternal gene expression values. The model fitting for each orthogroup was performed using this pruned species tree with the normalized TPM values at each tip. All models were fit on univariate data, i.e., single orthogroup with associated expression values or fold changes. A filtering step was inserted for the minimum tree size as some phylogenetic models are sensitive to sample size (Cooper et al., 2016). Standard errors for each species were included during the model fitting. The error terms were calculated using biological replicates for each species.

Model fitting was performed using the R packages OUwie (Beaulieu et al., 2012) and geiger (Pennell et al., 2014). From the latter white noise models were used as null models. These models do not contain phylogenetic signals within them, rather the data is best explained by a normal distribution without any covariance present. From OUwie multiple models were included in the analysis (Table 1).

**Table 1.** Table describing the analyzed evolutionary models with the parameters included in the model. OU1- Ornstein-Uhlenbeck model, BM – Brownian motion model, BMS- Multi-regime Brownian motion model, OUM – Multi-regime Ornstein-Uhlenbeck model, OUMV – Multi-regime Ornstein-Uhlenbeck model with variable σ^2^ parameters, OUMA - Multi-regim Ornstein-Uhlenbeck model with variable α parameters, OUMVA - Multi-regime Ornstein-Uhlenbeck model with variable α and σ ^2^ parameters

| Model | Regime | $\theta$ | $\alpha$ | $\sigma$ |
| --- | --- | --- | --- | --- |
| OU1 | Single regime | $\theta$ | $\alpha$ | $\sigma^2$ |
| BM1 | Single regime | - | - | $\sigma^2$ |
| OUM | Multiple regimes | $\theta_1, \theta_2, \theta_3$ | $\alpha$ | $\sigma^2$ |
| BMS | Multiple regimes | $\theta_1, \theta_2, \theta_3$ | - | $\sigma^2_1, \sigma^2_2, \sigma^2_3$ |
| OUMV | Multiple regimes | $\theta_1, \theta_2, \theta_3$ | $\alpha$ | $\sigma^2_1, \sigma^2_2, \sigma^2_3$ |
| OUMA | Multiple regimes | $\theta_1, \theta_2, \theta_3$ | $\alpha_1, \alpha_2, \alpha_3$ | $\sigma^2$ |
| OUMVA | Multiple regimes | $\theta_1, \theta_2, \theta_3$ | $\alpha_1, \alpha_2, \alpha_3$ | $\sigma^2_1, \sigma^2_2, \sigma^2_3$ |

All selected models were fitted to each pruned orthogroup expression or fold change data. The winning models were first selected using second-order Akaike Information Criterion (AICc). After selecting the best fitting model, a permutation test was performed. For this, the data points were randomly shuffled across the examined species tree and the winning model was fitted to it. All AICc values from the 250 rounds of permutation were extracted and tested against the original AICc using the Kolmogorov-Smirnov test to examine if the original AICc value could be explained by chance. Models for downstream analysis were selected if their AICc values exceeded 0.5 and the permutation test returned a significant result (p-value ≤ 0.05). Parameters from models sufficing these criteria were recovered and further examined.

In addition to testing the evolutionary dynamics of the RNA content in oocytes, we set out to explore the evolutionary patterns of the temporal dynamics during the MZT. We utilized the steps outlined above on the fold change results from differential gene expression analysis to do this. These fold change values were used directly for modeling as suggested previously (Dunn et al., 2013).

The power of maternal gene expression to classify reproductive modes was also queried. In order to explore this, we set out to use phylogenetic logistic models using phylolm (Tung Ho & Ané, 2014). A null hypothesis of constant dependent variables was used for the logistic models. The competing hypothesis was univariate maternal gene expression value. For each reproductive mode, a logistic model was built and tested resulting in 3 models for each orthogroup. Each model then could possibly classify the tested reproductive mode. Model selection was performed using both AICc values and the likelihood ratio test. For a model to be considered for downstream analysis it had to suffice the criteria of AICc values above 0.5, a significant improvement in the fit according to the likelihood ratio test and 4 or more species had to be present for the tested reproductive mode. Selected models were included for downstream analysis.

## 3. Results

### 3.1 Maternal gene expressions and 3’-UTR lengths vary across Metazoa

We have found that the proportion of the genomes being expressed in oocytes varied across species, on average 41% of all genes are expressed. A notable exception was T.transversa, which had 71% of all annotated genes in the oocytes. We have discretized the gene expressions into 4 categories based on the TPM values. Genes having TPM < 2 were considered not expressed as suggested previously (Wagner et al., 2013). A gene was considered as having a low expression value where 2 ≤ TPM < 100, a medium expression value where 100 ≤ TPM < 1000, and a high expression value where TPM ≥ 1000. A shared feature across all species is the lack of genes with high expression. Patterns for medium or low expression genes varied across species. In Hexapoda, maternal gene expressions are skewed towards medium expression categories. In contrast, echinoderm species show enrichment in low expression categories. Maternal genes undergoing degradation during MZT are abundant in medium expression categories. Additionally, in the degraded maternal gene set the high expression category profiles are more abundant (Figure 1). When inspecting the fold changes associated with the above-mentioned maternally expressed genes we saw a pattern emerge. After the MZT most maternal transcripts with low expression values are downregulated. In contrast, the maternal transcripts with higher initial values show an upregulation after the MZT (Figure 1). Enrichment analysis of maternal genes revealed terms such as RNA splicing, mRNA transport, histone acetylation, and transcription coactivator activity were significantly enriched across species. We found that such features were commonly enriched across most species. For maternal genes which undergo clearance during MZT, our analysis revealed less overlapping enriched terms (Suppl. figure 4).

**Figure 1.**
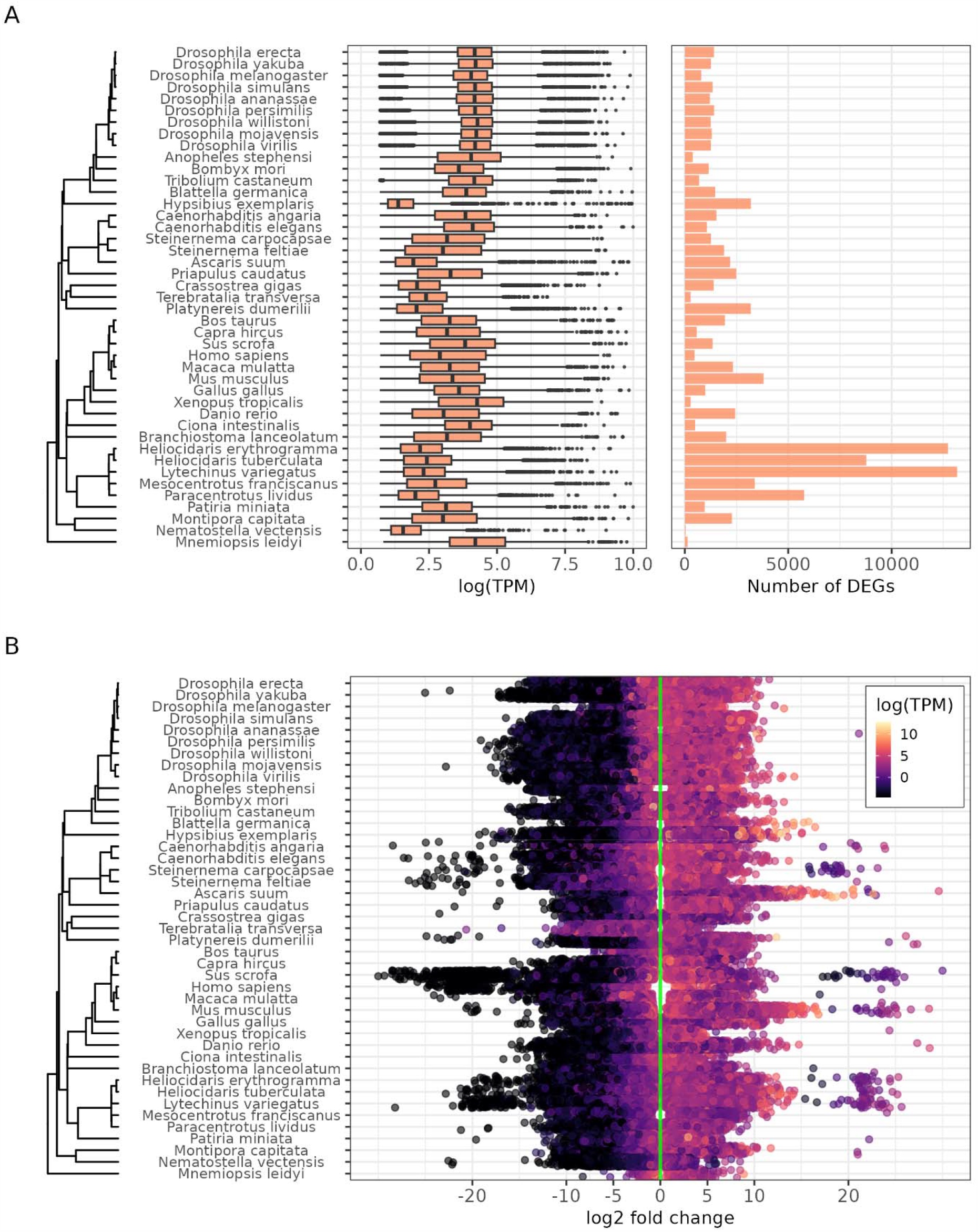
Maternal gene expression and fold change patterns across the metazoan tree. (A) Distribution of maternal gene expressions across the analyzed species on the left and number of degraded maternal genes for each species on the right. (B) Fold change patterns of maternal genes across the studied species. Genes with lower expression values tend to be downregulated across all included species.

The length of 3’-UTRs varies between different species, with ecdysozoan species generally having shorter 3’-UTRs compared to deuterostome species. However, *C. hircus* is an exception with longer 3’-UTR sequences. In general, non-downregulated and downregulated genes within a species have longer 3’-UTR sequences compared to other genes without a maternal expression, which may suggest that post-transcriptional regulation plays a greater role in these genes as previously suggested (Mishima & Tomari, 2016). However, exceptions exist in *H. sapiens* and *S. carpocapsae*, which have equal 3’-UTR lengths between maternally expressed and not expressed genes. Additionally, hexapod species show shorter 3’-UTR sequences in non-downregulated genes compared to downregulated genes, indicating that the stability of maternal transcripts may depend more on 3’-UTR sequences in this clade. When examining the coding sequence (CDS) lengths the non-downregulated and downregulated genes were found to have longer CDS (Suppl. figure 19).

### 3.2 Phylogenetic signal is present in the maternal gene expression dataset and justifies the use of evolutionary models

Our analysis on the expression data where orthogroups met the species tree cutoff revealed a phylogenetic signal present in 57% of cases, justifying the use of phylogenetic comparative methods. The K statistic estimates showed a majority of orthogroups with higher dissimilarity than expected (K < 1), while a smaller portion suggested more homogeneous expressions (K > 1). Similar results were obtained for the fold change data, with a phylogenetic signal present in 68% of cases and a left-skewed distribution of the K statistic (Suppl. figure 5).

### 3.3 Both selection and neutral drift is present in maternal gene expression evolution

Evolutionary models have been fitted to maternal gene expression datasets. Both gene expression and fold change data have been included. In total, for each orthogroup 8 model fits were tested simultaneously and the best-fitting model was selected based on both AICc weights and permutation tests. The AICc weight distribution for each winning model varied. For some (BM1 and OU1) it was symmetrical at around 0.5, whilst for others it was skewed to the right (OUM, OUMA, OUMV, OUMVA, BMS) (Suppl. figure 6). Furthermore, no apparent differences are noticeable between the fits of expression values and fold change values as they have highly similar distributions. This implies that more complicated models have better fitting models, reflecting a complicated evolutionary landscape when it comes to the expression of maternal genes.

After filtering AICc weights and the significant permutation tests 3825 orthogroups remained for downstream analyses. Out of these 1105 were represented in both expression datasets and fold change datasets (Figure 2). The most abundant models for maternal gene expressions were the OU models with single optima and Brownian motion models with multiple σ^2^ values. The same was true for the fold changes between oocytes and embryos right after MZT. More complicated models had been recovered more sparsely. Generally, the more parameters to be estimated the less likely the model would win. Interestingly, the Brownian-motion model with a single σ^2^ model was the least likely to win in the fold change dataset, whilst its more complicated version was the most likely to win. Most of the winning models were also associated with a significant phylogenetic signal (Suppl. figure 6). Models, where not all orthogroups had significant phylogenetic signals, were mostly OU models and the Brownian motion model with multiple σ^2^, which is not surprising as Blomberg’s K test assumes a Brownian motion generating the data (Blomberg et al., 2003). Furthermore, the K-statistic, which provides an indication for the departure from the expected distribution of Brownian motion showed that BM1 models were the ones closest to the expected distribution. More complicated models showed a greater departure from the expected distribution. Orthogroups in the different models were enriched for various gene ontological terms with overlap across the models. Shared functions across some models were centered around splicing events, mRNA processing, and mitochondrial processes. OU1 models were uniquely involved in processes involved in DNA replication and vesicular transport. Uniquely enriched processes in other models were that of transcription and translation regulation of BMS models (Suppl. figure 8).

**Figure 2.**
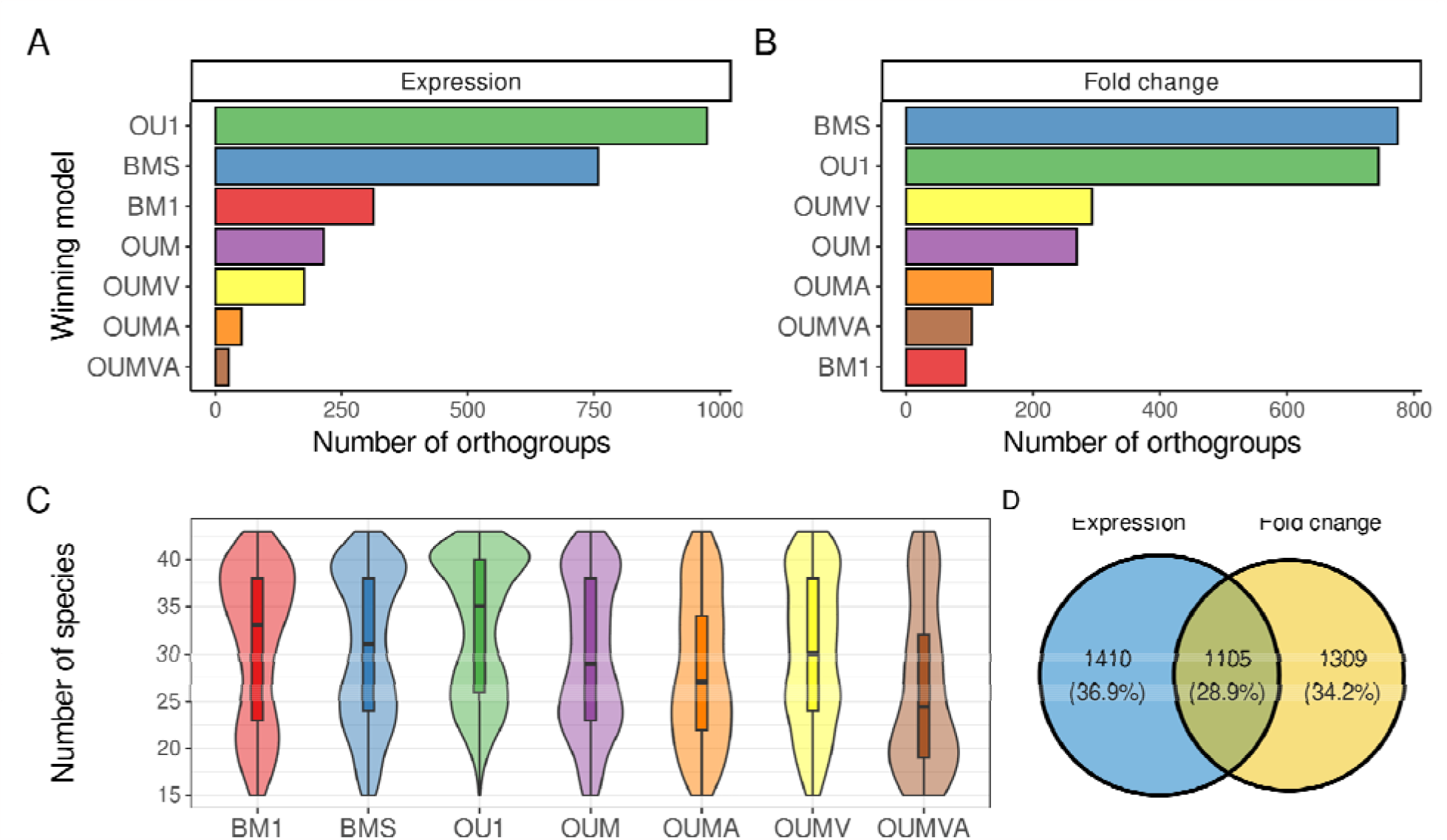
The best fitting evolutionary models. (A) The number of orthogroups belonging to each tested evolutionary model in the case of the expression dataset. Only best fitting models (AICc ≥ 0.5 and permutation test p-value ≤ 0.05) are represented. (B) The same as panel (A) with the fold change dataset used. (C) The distribution of the species present in each orthogroups after pruning for each evolutionary model tested. (D) Venn diagram of the number of orthogroups included in downstream analyses (after filtering for AICc and permutation test). In blue are the number of orthogroups for the expression dataset and in yellow are the number of orthogroups for the fold change dataset. The overlap includes orthogroups where both the expression and the fold change dataset has a best fitting significant evolutionary model present.

### 3.4 Parameter estimates of evolutionary models reflect evolutionary patterns across reproductive modes

The ⍰ estimates from the OUM model fittings showed higher variances for hemotrophic viviparity compared to the two other reproductive modes (Figure 3). ⍰ estimates for the expression values are of negligible effect size (Suppl. table 3). Oviparous species show a slight bias towards ⍰ estimates above 0 for fold change datasets. This bias is not present in ovuliparous species, as here the estimated ⍰ values show a slight bias for values below 0. Most frequently the highest optima were found to be in hemotrophic viviparity for both expression values and fold changes. Noticeably different was the least frequent highest optima for ovuliparous species. For the lowest optima in each orthogroup, a different pattern emerged. Yet again, ovuliparous species rose to the top when it comes to the lowest number of maternal transcripts found in these species. Additionally, ovuliparous species had the most frequently lowest optima of fold changes. The opposite of this was the hemotrophic viviparous species, which had the least frequently lowest optima.

**Figure 3.**
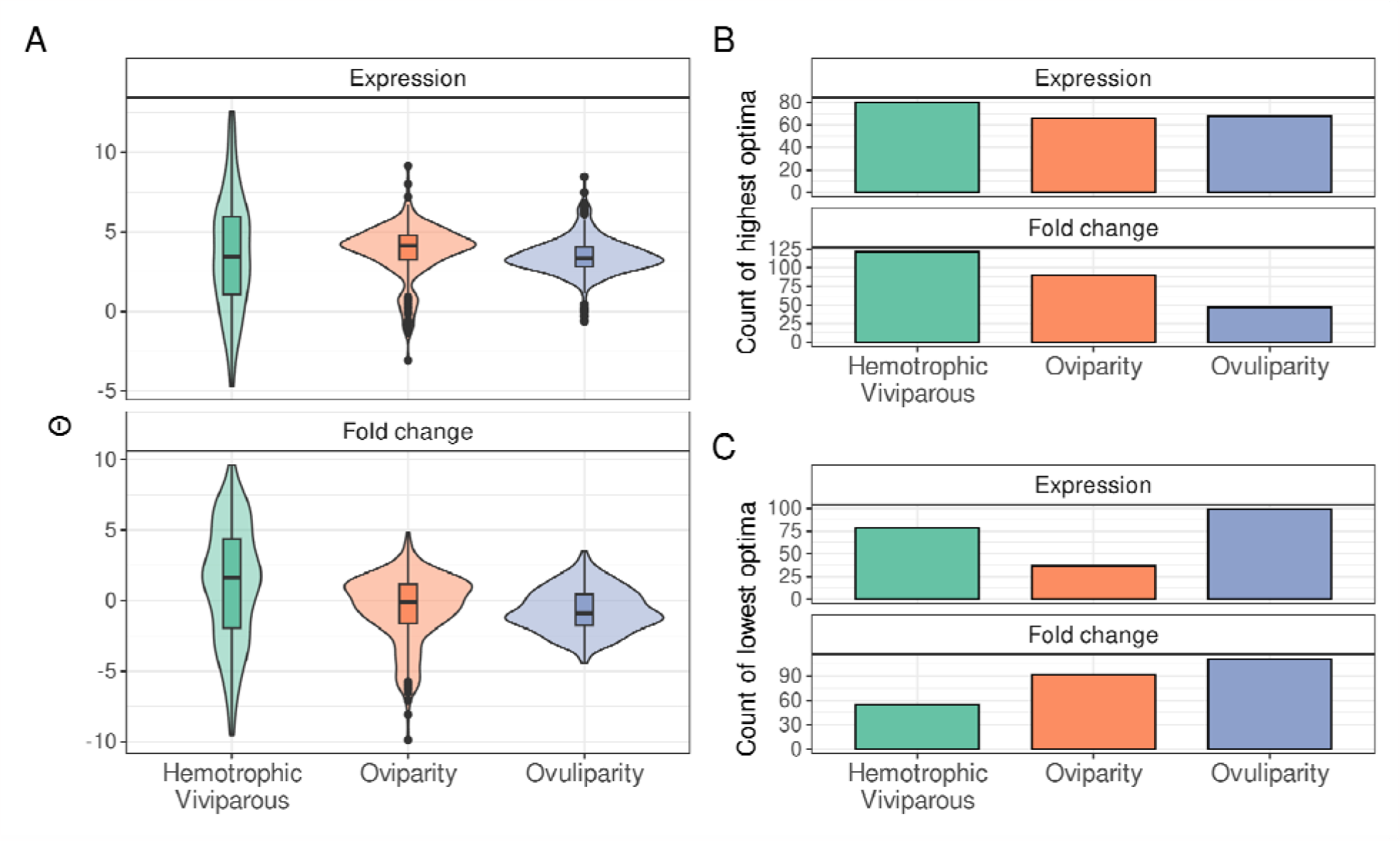
⍰ estimates for orthogroups where OUM models were found to be the best fitting model. (A) Distribution of ⍰ estimates for the reproductive modes in both expression dataset (above) and fold change dataset (below). (B) The number of highest ⍰ values for both datasets in all three reproductive modes. (C) The number of lowest ⍰ values for both datasets in all three reproductive modes.

In the case of orthogroups where α was allowed to vary across regimes generally the optima were similar, except for ovuliparous species expression values, which were slightly lower (with moderate effect size). No sizable differences (i.e. small or negligible Cohen’s D values) were observable between the α values across regimes (Suppl. table 4). The highest ⍰ for orthogroups with OUMA models were abundant in ovuliparous species most commonly (Suppl. figure 10). Contrary to this, the lowest ⍰ characterized ovuliparous species. Supporting this pattern were the α value estimates, where the highest α values generally characterized oviparous species best. Contrary to the lowest ⍰ estimates, the lowest α values displayed a more obscure pattern. Here no strong differences were noticeable for expression values, whilst ovuliparous species displayed the lowest α values overall for OUMA models. Similarly to the variable alpha models, the variable σ^2^ models did not show a strong difference between their ⍰, nor between the σ^2^ values (Suppl. figure 11, Suppl. table 5). Oviparous species displayed the most frequent highest ⍰ for expression values, whilst hemotrophic viviparous species most frequently had the highest optima in the fold change datasets. The lowest optima for both expression values and fold changes were most common in hemotrophic viviparous mode. Interestingly a strong signal was noticeable for the varying σ^2^ estimates. These were most frequently the lowest for oviparous species and least frequently highest. A big proportion of the winning models followed a Brownian motion model with variable σ^2^ values. Here, for the parameter estimates, we noticed that the 1 value estimates were highly variable (Suppl. figure 12). Interestingly, the fitted models returned realistic estimates for the ancestral state of the ⍰ values. The σ^2^ values displayed differences of large effect sizes (Suppl. table 6), most notably hemotrophic viviparous species had high σ^2^ estimates which were most frequently the highest σ^2^ values per orthogroup. Oviparous species had the lowest σ^2^ estimates in their expression values.

Parameter estimates in the case of OU1 and BM1 models showed correlated estimates (Suppl. figure 13). σ^2^ and α parameters in the case of OU1 models were strongly correlated, as were the mean expression values and the ⍰ estimates for both OU1 and BM1 models. Interestingly the σ^2^ and α parameters negatively correlated with both the species included in the model fitting process and the mean expression values. A positive correlation was observable between mean expression values and the count of species included during model fitting. The positive correlation between mean expression and species numbers was observable across all included models, although at varying degrees (Suppl. figure 13, Suppl. figure 14, Suppl. figure 15, Suppl. figure 16, Suppl. figure 17). The same holds true for α and σ^2^ parameters. Interestingly, in the OUM models the correlations between ⍰ and mean expression are spurious. For fold changes, some of these associations are completely lost and no linear relationship is present between optima and mean expression values. Only weak or no correlations are present between the parameters of BMS models (Suppl. figure 17). A notable exception to this is the strong negative association between ⍰ of the oviparous species and ovuliparous species. Correlation coefficients between the OUMA and OUMV model parameters followed the trend outlined above. In the OUMA models the α parameters from the oviparous species do not correlate significantly with either ovuliparous or hemotrophic viviparous α parameters, nor do they significantly correlate with the σ^2^ parameters. A similar pattern emerged in the case of OUMV models.

### 3.5 Gene expression and fold change data are sufficient to distinguish reproductive modes from each other

Phylogenetic logistic models were successful in classifying reproductive modes based on either maternal gene expression data or fold changes of maternal genes during MZT. Some data points contained enough signal to distinguish reproductive modes from each other (Figure 4.). Likelihood ratio tests and AICc weights were used to select plausible models. In total, for gene expression data 47 models and for fold change data 157 models were selected. For models distinguishing ovuliparous species terms involved in ribosomal function of cytoskeletal functions were enriched. Enriched terms were found for fold change data points distinguishing hemotrophic viviparous species also. Terms such as nervous system development, hair follicle development or functions regarding binding activities were enriched. Apart from the previous two enrichment results, no further significant enrichments were found.

**Figure 4.**
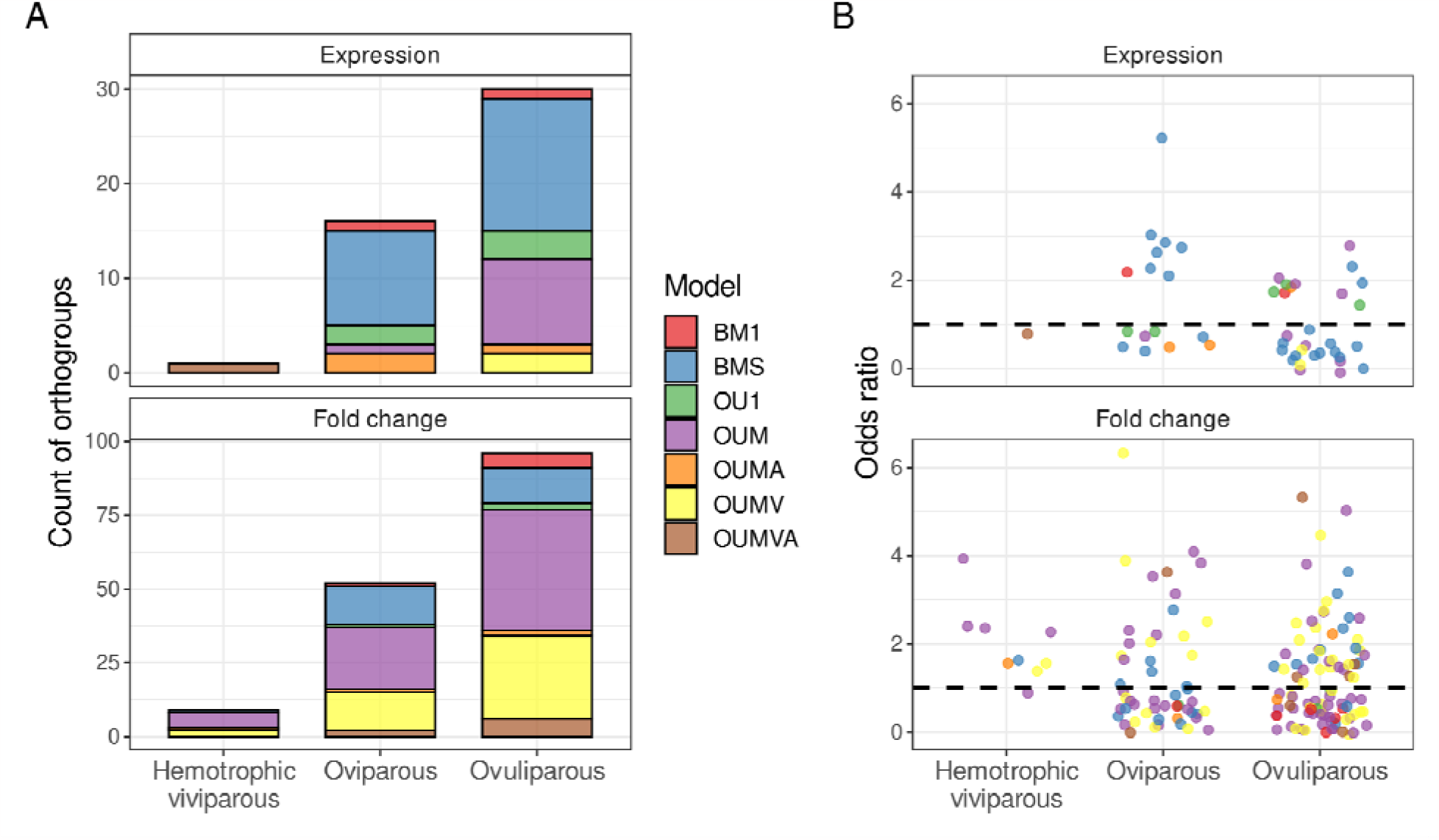
Summary of phylogenetic logistic modeling. (A) Number of orthogroups with sufficient phylogenetic signal present t classify reproductive mode. (B) Odds ratios of phylogenetic logistic models. Dashed lines represent odds ratio of 1. Values above the line suggest that data values for that reproductive mode are higher than the other reproductive modes. Values below the dashed line suggest that data values for that reproductive mode are lower than the other reproductive modes.

Overall, the ovuliparous reproductive mode is the most frequently classifiable, and the hemotrophic viviparous reproductive mode is the least. There is also a bias in the evolutionary models of the orthogroups for each mode. For gene expression data points the BMS models are relatively the most abundant, whilst for fold change data the OUM and OUMV models are the most prevalent. The logistic model odds ratios do not suggest that any phylogenetic model would contribute more likely to the classification of reproductive modes.

Within the same orthogroup a logistic model could distinguish between multiple reproductive modes. Such is the case for the orthogroup annotated as ribonuclease H1, where two separate models emerged. In one of the models, the ovuliparous species could be distinguished by the upregulation of the ribonuclease, whilst for the oviparous species the exact opposite was noticeable.

## 4. Discussion

Our findings reveal several observations that shed light on maternal gene expression during early development. Furthermore, we found evidence supporting a more intricate evolutionary pattern of maternal gene expression, compared to what was previously described (Atallah & Lott, 2018b, 2018a; Cruickshank & Wade, 2008; Demuth & Wade, 2007; Mousseau & Fox, 1998).

### 4.1 Potential factors influencing MZT dynamics

Our assimilated datasets showed that maternal genes possess a variable expression pattern across species and clades. An apparent bias is generally present toward weakly or moderately expressed maternal genes with a big proportion of the genome. Additionally, the weakly expressed maternal genes are more likely to be degraded after the MZT, whilst moderately expressed maternal genes show a tendency towards being upregulated. This upregulation could be due to two factors. Firstly, it could originate from *de novo* transcription from the zygotic genome. Secondly, it could have its roots in the polyadenylation of the transcripts and thereby making such transcripts visible during polyA tail selection. However, the data assimilated in this study is not adequate to distinguish between such origins. Nevertheless, both processes result in an increase in the availability of transcripts for translation, suggesting an important role for such transcripts during and right after MZT. Our results provide further evidence that maternal genes during MZT are regulated through their 3’-UTR sequences across a wide array of clades in the Metazoan tree.

### 4.2 Evolutionary model fittings and departure from Brownian motion

Our evolutionary model fittings revealed a significant presence of phylogenetic signal in the expression data. This prompted further investigation into evolutionary modeling as the Blomberg’s K values suggested a departure from the assumption of the expected Brownian motion. We competed multiple evolutionary scenarios for each maternal gene expression data and fold change data and set out to find the best explanation for them. The assumption based on previous studies (Cruickshank & Wade, 2008; Demuth & Wade, 2007; Kirkpatrick & Lande, 1989; Mousseau & Fox, 1998) was a great deal of divergence, reflected in Brownian motion models. Some, more recent publications on drosphiliid species suggest a more dynamic landscape of maternal gene expression evolution (Atallah & Lott, 2018b, 2018a). We expanded these works with a broader sampling and testing embedded in a phylogenetic framework. Our results contradict the expected divergence, rather they align more with the more recent results of a dynamic evolutionary landscape. A possible explanation for the discrepancy between nucleotide data and current results based on expression data could be attributed to the pleiotropic effects of nucleotide changes (Paaby & Rockman, 2013). This effect could be alleviated altering the expression levels of maternal genes, as this scenario would enable plasticity without affecting later developmental stages. Divergence could be achieved by varying the levels of expressions of genes, thereby introducing new possibly advantageous molecular functions in the maternal gene pool. This notion is supported by our finding that BMS and BM1 models are well represented in the expression datasets. In parallel to such divergences, a selective force maintains the expression for a set of maternal genes essential for basic cellular functions and general molecular mechanisms during early development. This is exemplified by the prevalence of OU1 models and multi-regime OU models.

### 4.3 Evolutionary scenarios for maternal gene expression and fold change data

Including multi-regime models in our analysis enabled us to inspect differences across different reproductive modes. OUM models provided many insights into such evolutionary differences across reproductive modes. The relatively bigger spread of ⍰ estimates for hemotrophic viviparous species across all multi-regime OU models suggested a scenario where due to the stable environment of the placenta variation in optima is permissible. Fold change data for such species also supports this notion, as a relatively bigger spread was present for the ⍰ estimates. A slight bias towards upregulation in hemotrophic viviparous species suggests a tendency to favor the upregulation of maternal genes in such species for orthogroups following OUM models. Alternatively, spurious estimates could be explained by a sampling bias. Hemotrophic viviparous species are underrepresented in our dataset, leading to potentially spurious parameter estimates. Sampling bias could also be held accountable in OUMA and OUMV α and σ^2^ estimates. The overall emerging pattern for orthogroups with such models is that oviparous species have higher α and lower σ^2^ parameters, suggesting selective forces are more prevalent in maintaining expression values and fold changes in oviparous species. The bias could arise in the relative oversampling of drosophilid lineages with relatively less change among them compared to other lineages with fewer representatives and result in estimates reflecting this. A similar scenario could be present in orthogroups following BMS models, here the σ^2^ estimates are highest for hemotrophic viviparous species, which could arise from the spurious sampling mentioned above.

### 4.4 Evolutionary differences across reproductive modes

Oviparous species also display a slight tendency towards upregulation during MZT, whilst ovuliparous species display a slight preference towards downregulation. This suggests that for ovuliparous embryos more maternal genes are downregulated, despite being present in the maternal transcriptome, compared to other modes of reproduction. This notion is further supported by our results showing that ovuliparous species have the lowest optima estimated most frequently. A possible explanation for this observation could be that the progeny of such species is highly susceptible to environmental stressors. Selection could favor a scenario where there is less constraint on what genes are present in the maternal transcriptome. This could occur as means of buffering the transcriptional quiescence while still enabling plasticity towards environmental stressors in combination with ovuliparous species having zygotic transcription initiated relatively later during embryogenesis (Schulz & Harrison, 2019). Such a scenario would then suggest that in a stable environment, a specific set of maternal genes is provided by the maternal organism as transcription is costly (Wang et al., 2015). We found evidence backing this suggestion up for hemotrophic viviparous species, where the optima were the highest in most cases for expression values and upregulation of maternal genes was more likely than downregulation.

Correlation structures across parameter estimates, mean values of data, and pruned tree sizes added further details to the scenario outlined above. Across all models, the estimates suggested to varying degrees that there is a linear relationship between the pruned tree size and the mean values of data. This linear relationship was the strongest and most positive in single-regime models, meaning as more species have maternal expression associated with orthogroups, the stronger its expression values. A sensible explanation would then be if a given gene is universally required for early embryogenesis selection would favor that gene being more strongly expressed and across more species. This association is dissipated in some orthogroups following extended OU models, which further strengthens the notion outlined above. In multi-regime models not all regimes have the same selective regimes present, therefore the association weakens compared to scenarios where all regimes evolve under similar circumstances. The overall positive correlation between means of expression and fold change optima estimates across all multi-regime OU models further strengthens the conclusion drawn above, whereby genes with low expression show a tendency towards being downregulated during MZT. The strong overall positive correlation between α and σ^2^ could be potentially traced back to the sampling biases and restrictions outlined above. This is further supported by the weak negative association between pruned tree sizes, i.e. data points, and parameter estimates.

### 4.5 Implications of gene expression differences in reproductive modes

Our analyses also provided a set of genes with sufficient signal to distinguish between reproductive modes. This could have several implications. Firstly, it suggests that maternal gene expressions and developmental dynamics are influenced by the reproductive mode. Secondly, it expands further our current knowledge on the biology of oocytes and early developmental steps across species. Our result that ribonuclease H1 is upregulated in ovuliparous species and downregulated in oviparous species depicts a scenario where for species with both reproductive modes the clearance of RNAs is pivotal through ribonucleases. The origin of such ribonucleases is different, for ovuliparous species it could originate from either *de novo* transcription or polyadenylation of the transcripts, for oviparous species ribonuclease transcripts are *a priori* expressed before the MZT. Our results are limited by the scarce sampling of species. With more species sampled, we could get a finer resolution of which maternal genes are pivotal for which reproductive modes. Furthermore, these results suggest that such classifiers with more species under investigation have the potential to identify pivotal maternal genes for a *priori* defined clades. This approach could enable the discovery of critical maternal genes involved in primate reproduction and therefore of relevance for human reproductive health.

### 4.6 Proposed hypothesis for the evolutionary landscape of maternal gene expression

Based on our results we propose a hypothesis for the evolutionary landscape of maternal gene expression evolution. A core set of maternal genes with functions necessary for early divisions and initiation of development are under selective constraint to be expressed across a multitude of species. This is in line with the universal requirement of cellular functions across all species during early development. This core group of maternal genes has its temporary dynamics also under constraint during MZT. Apart from this core set of maternal genes, a set of variable maternal genes could also be identified. This set varies across species and shows no constraint on its expression values. Compared to the core set of maternal genes, the variable set can be characterized by weak and variable expression, suggesting a stochastic process behind it, for example, leaky transcription. To better adapt to unpredictable environmental stressors experienced by early embryos in ovuliparous and oviparous species, it may be advantageous to produce *a priori* genes in a stochastic manner that can respond to those specific stressors. Therefore, selection would maintain the presence of such variably expressed maternal genes as they could still offer a selective advantage.

